# Modern wolves trace their origin to a late Pleistocene expansion from Beringia

**DOI:** 10.1101/370122

**Authors:** Liisa Loog, Olaf Thalmann, Mikkel-Holger S. Sinding, Verena J. Schuenemann, Angela Perri, Mietje Germonpré, Herve Bocherens, Kelsey E. Witt, Jose A. Samaniego Castruita, Marcela S. Velasco, Inge K. C. Lundstrøm, Nathan Wales, Gontran Sonet, Laurent Frantz, Hannes Schroeder, Jane Budd, Elodie-Laure Jimenez, Sergey Fedorov, Boris Gasparyan, Andrew W. Kandel, Martina Lázničková-Galetová, Hannes Napierala, Hans-Peter Uerpmann, Pavel A. Nikolskiy, Elena Y. Pavlova, Vladimir V. Pitulko, Karl-Heinz Herzig, Ripan S. Malhi, Eske Willerslev, Anders J. Hansen, Keith Dobney, M. Thomas P. Gilbert, Johannes Krause, Greger Larson, Anders Eriksson, Andrea Manica

**Affiliations:** Palaeogenomics & Bio-Archaeology Research Network Research Laboratory for Archaeology and History of Art, University of Oxford, Dyson Perrins Building, South Parks Road, Oxford OX1 3QY, UK; Department of Zoology, University of Cambridge, Downing Street, Cambridge CB2 3EJ, UK; Manchester Institute of Biotechnology, School of Earth and Environmental Sciences, University of Manchester, Manchester, M1 7DN, UK; Department of Pediatric Gastroenterology and Metabolic Diseases, Poznan University of Medical Sciences, Szpitalna 27/33, 60-572 Poznan, Poland; Centre for GeoGenetics, Natural History Museum of Denmark, University of Copenhagen, Øster Voldgade 5-7, DK-1350 Copenhagen, Denmark; Natural History Museum, University of Oslo, P.O. Box 1172 Blindern, NO-0318 Oslo, Norway; The Qimmeq project, University of Greenland, Manutooq 1, PO Box 1061, 3905 Nuussuaq, Greenland; Institute for Archaeological Sciences, University of Tübingen, Rümelinstr. 23, 72070 Tübingen, Germany; Senckenberg Centre for Human Evolution and Palaeoenvironment, University of Tübingen, 72070 Tübingen, Germany; Department of Human Evolution, Max Planck Institute for Evolutionary Anthropology, Deutscher Platz 6, 04103 Leipzig, Germany; OD Earth and History of Life, Royal Belgian Institute of Natural Sciences, Vautierstraat 29, 1000 Brussels, Belgium; Department of Geosciences, Palaeobiology, University of Tübingen, Tübingen, Germany; School of Integrative Biology, University of Illinois at Urbana-Champaign, 109A Davenport Hall, 607 S. Mathews Avenue, Urbana IL 61801, USA; OD Taxonomy and Phylogeny, Royal Belgian Institute of Natural Sciences, Vautierstraat 29, 1000 Brussels, Belgium; Faculty of Archaeology, Leiden University, Postbus 9514, 2300 RA Leiden, The Netherlands; Breeding Centre for Endangered Arabian Wildlife, PO Box 29922 Sharjah, United Arab Emirates; Mammoth Museum, Institute of Applied Ecology of the North of the North-Eastern Federal University, ul. Kulakovskogo 48, 677980 Yakutsk, Russia; National Academy of Sciences, Institute of Archaeology and Ethnography, Charents St. 15, Yerevan 0025, Armenia; Heidelberg Academy of Sciences and Humanities: The Role of Culture in Early Expansions of Humans, Rümelinstr. 23, 72070 Tübingen, Germany; Departement of Anthropology, University of West Bohemia, Sedláčkova 15, 306 14 Pilzen, Czech republic; Moravian museum, Zelný trh 6, 659 37 Brno, Czech republic; Hrdlička Museum of Man, Faculty of Science, Charles University, Viničná 1594/7,128 00 Praha, Czech republic; Institute of Palaeoanatomy, Domestication Research and History of Veterinary Medicine, Ludwig-Maximilians-University Munich, Kaulbachstraße 37 III/313, D-80539 Munich, Germany; Geological Institute, Russian Academy of Sciences, 7 Pyzhevsky per., 119017 Moscow, Russia; Institute for Material Culture History, Russian Academy of Sciences, 18 Dvortsovaya nab., St Petersburg 191186, Russia; Arctic and Antarctic Research Institute, 38 Bering St., St Petersburg 199397, Russia; Insitute of Biomedicine and Biocenter of Oulu, Medical Research Center and University Hospital, University of Oulu, Aapistie 5, 90220 Oulu University, Finland; Carl R. Woese Institute for Genomic Biology, University of Illinois at Urbana-Champaign, 1206 W Gregory Dr., Urbana, Illinois 61820, USA; Wellcome Trust Sanger Institute, Hinxton, Cambridge CB10 1SA, UK; Department of Archaeology, Classics and Egyptology, University of Liverpool, 12-14 Abercromby Square, Liverpool L69 7WZ, UK; Department of Archaeology, University of Aberdeen, St Mary’s, Elphinstone Road, Aberdeen AB24 3UF, UK; Department of Archaeology, Simon Fraser University, Burnaby, B.C. V5A 1S6, 778-782-419, Canada; Norwegian University of Science and Technology, University Museum, N-7491 Trondheim, Norway; Max Planck Institute for the Science of Human History, Khalaische Straße 10, 07745 Jena, Germany; Department of Medical & Molecular Genetics, King’s College London, Guys Hospital, London SE1 9RT, UK

## Abstract

Grey wolves (*Canis lupus*) are one of the few large terrestrial carnivores that maintained a wide geographic distribution across the Northern Hemisphere throughout the Pleistocene and Holocene. Recent genetic studies have suggested that, despite this continuous presence, major demographic changes occurred in wolf populations between the late Pleistocene and early Holocene, and that extant wolves trace their ancestry to a single late Pleistocene population. Both the geographic origin of this ancestral population and how it became widespread remain a mystery. Here we analyzed a large dataset of novel modern and ancient mitochondrial wolf genomes, spanning the last 50,000 years, using a spatially and temporally explicit modeling framework to show that contemporary wolf populations across the globe trace their ancestry to an expansion from Beringia at the end of the Last Glacial Maximum - a process most likely driven by the significant ecological changes that occurred across the Northern Hemisphere during this period. This study provides direct ancient genetic evidence that long-range migration has played an important role in the population history of a large carnivore and provides an insight into how wolves survived the wave of megafaunal extinctions at the end of the last glaciation. Moreover, because late Pleistocene grey wolves were the likely source from which all modern dogs trace their origins, the demographic history described in this study has fundamental implications for understanding the geographical origin of the dog.

---

The Pleistocene epoch harbored a large diversity of top predators though most became extinct during or soon after the Last Glacial Maximum (LGM) ~24 thousand years ago. The grey wolf (*Canis lupus*) was one of the few large carnivores that survived and maintained a wide geographical range throughout the period (1), and both the paleontological and archaeological records attest to the continuous presence of grey wolves across the Northern Hemisphere for at least the last 300,000 years (2) (reviewed in Supplementary Information 1). This geographical and temporal continuity across the Northern Hemisphere contrasts with analyses of complete modern genomes which have suggested that all contemporary wolves and dogs descend from a common ancestral population that existed as recently as ~20,000 years ago (3–5). These analyses point to a bottleneck followed by a rapid radiation from an ancestral population around or just after the LGM, but the geographic origin and dynamics of this radiation remain unknown. Resolving these demographic changes is necessary for understanding the ecological circumstances that allowed wolves to survive the late Pleistocene megafaunal extinctions. Furthermore, because dogs were domesticated from late Pleistocene grey wolves (6), a detailed insight into wolf demography during this time period would provide an essential context for reconstructing the history of dog domestication.

Reconstructing past demographic events solely from modern genomes is challenging since multiple demographic histories can lead to similar genetic patterns in present-day samples (7). Analyses that incorporate ancient DNA sequences can eliminate some of these alternative histories by quantifying changes in population genetic differences through time. While nuclear markers provide greater power relative to mitochondrial DNA (mtDNA), the latter is more easily retrievable and better preserved in ancient samples due to its higher copy number compared to the nuclear DNA, thus allowing for the generation of datasets with greater geographical and temporal coverage. Furthermore, the nuclear mutation rate in canids is poorly understood, leading to wide date ranges for past demographic events reconstructed from panels of modern whole genomes (e.g. (3, 5). Having samples from a broad time period can reduce mutation rate uncertainty by calibrating the evolutionary rate with directly dated samples (8–10).

Although mtDNA can be retrieved from a wider range of ancient samples, the sparseness of samples across space and time, compounded by the stochasticity of a single non-recombining genetic marker, can lead to patterns that are difficult to interpret intuitively (7). Population genetic models that explicitly capture the expected temporal (e.g. (11) and spatial (e.g. (12, 13) differences between samples can be used to overcome this problem.

In order to reconstruct the demographic history of wolves, we assembled a substantial dataset (Fig. 1, Table S1) consisting of 90 modern and 45 ancient wolf whole mitochondrial genomes (55 of which are newly sequenced), spanning the last 50,000 years and the geographic breadth of the Northern Hemisphere. We first used the ancient mitogenomes to estimate a wolf mutation rate using BEAST. We then designed a spatially and temporally explicit population genetic (coalescent) model that accounts for the stochasticity of the mitochondrial phylogenetic tree, as well as the uneven spatial and temporal distribution of the samples. We used our spatial model to investigate the origin and population dynamics of the expansion of grey wolves during the LGM.

**Fig. 1.**
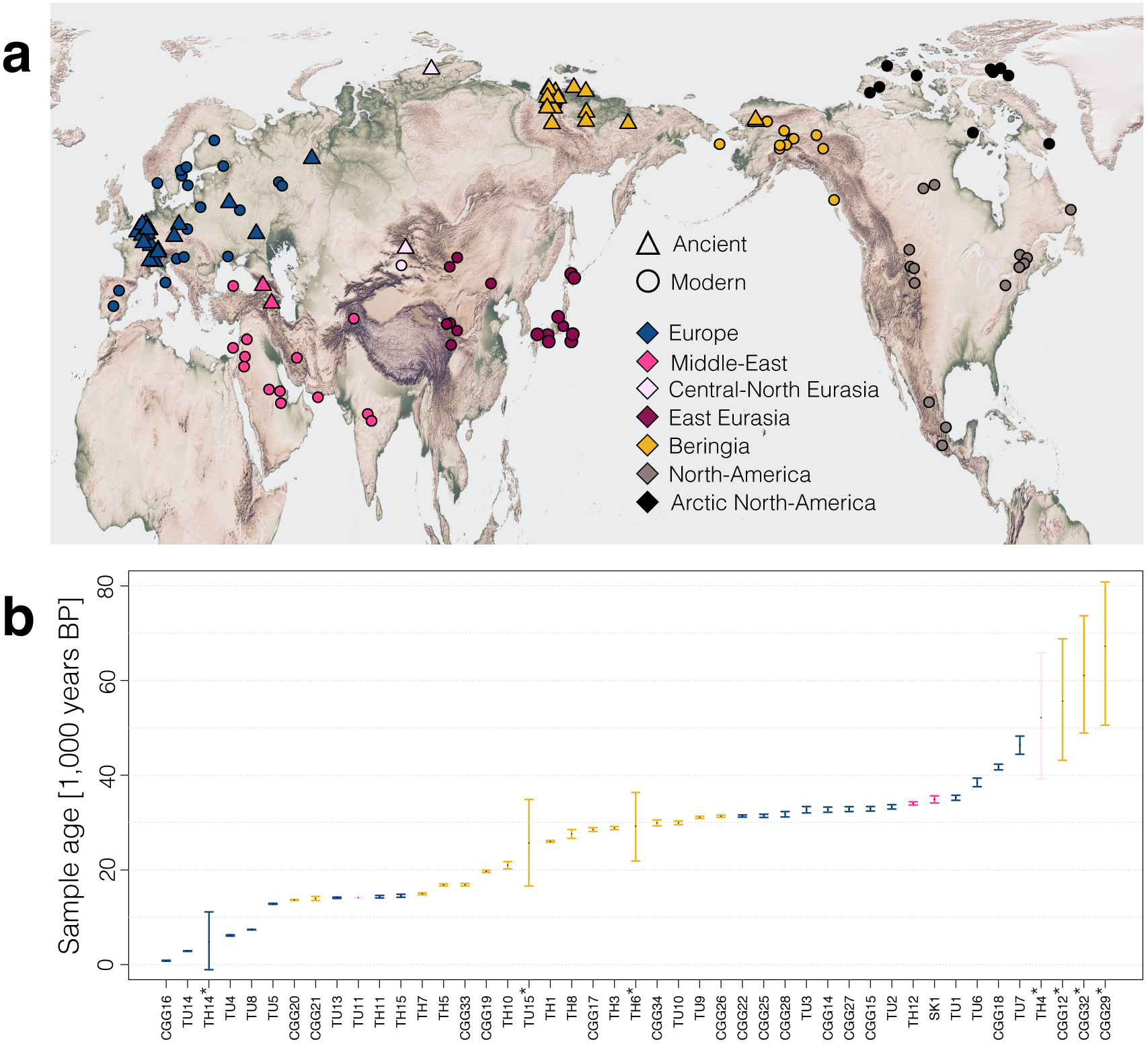
Geographic distribution of modern (circles) and ancient (triangles) samples, grouped into seven geographic regions (demes, colour coded) (a) and temporal distribution of ancient samples (b) used in the analyses. * Samples dated by molecular dating.

## Bayesian Phylogenetic Analysis

All sequences included in the study were subjected to stringent quality criteria with respect to coverage and damage patterns. We used the 38 ancient samples for which we had direct radiocarbon dates to estimate mitochondrial mutation rates using BEAST (14), and molecularly dated the remaining seven ancient samples (Supplementary Information 3.1).

Our Bayesian phylogenetic analysis suggests that the most recent common ancestor of all North Eurasian and American wolf samples dates to ca. 90,000 (95% CIs: 82.000 – 99.000) years ago (Fig. 2, see Figs. S11 and S12 for node support values and credibility intervals). At the root of this tree, we find a divergent clade consisting exclusively of ancient samples from Europe and the Middle East that has not contributed to present day mitochondrial diversity in our data (see also (15). The rest of the tree consists of a monophyletic clade made up of ancient and modern samples from across the Northern Hemisphere and shows a pattern of rapid bifurcations of genetic lineages centered on 25,000 years ago. A Bayesian skyline analysis (Fig. S13, see Supplementary Information 3 for details) also shows a recent reduction in effective population size. This pattern is compatible with a scenario of rapid radiation that has also been suggested by whole genome studies (e.g. (3, 5). At the root of this clade we find predominantly samples from Beringia, pointing to a possible expansion out Northeast Eurasia or the Americas. However, given the uneven temporal and geographic distribution of our samples and the stochasticity of a single genetic marker (16), it is important to explicitly test the extent to which this pattern can occur by chance under other plausible demographic scenarios, taking the geographic and temporal distribution of our samples into account.

**Fig. 2.**
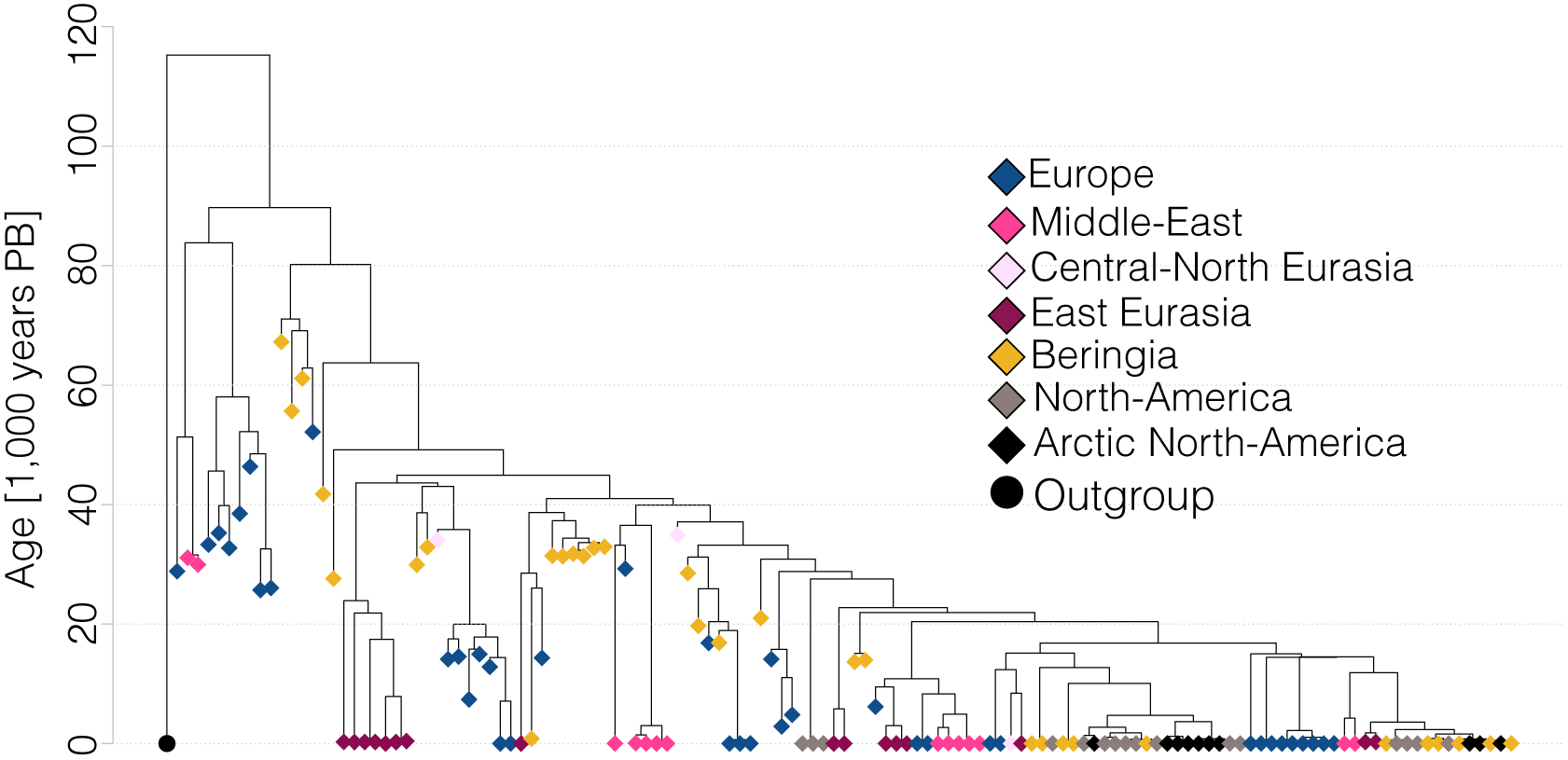
Tip calibrated BEAST tree of all samples used in the spatial analyses (diamonds), coloured by geographic region. The circle represents an outgroup (modern Indian wolf, not used in the analyses).

## Spatial Modelling of Past Wolf Demography

Motivated by the population structure observed in whole genome studies of modern wolves (5), we tested the degree of spatial genetic structure among the modern wolf samples in our dataset, and found a strong pattern of genetic isolation by distance across Eurasia (*ρ*=0.3, *p*<0.0001; see Fig. S8). To take this structure into account in our spatial framework, we represented the wolf distribution in the Northern Hemisphere as seven demes (Fig. 1), each of which is defined by major geographic barriers including mountain ranges, seas, oceans and deserts (see Materials and Methods). We then used coalescent simulations (see Materials and Methods) to test a range of different explicit demographic scenarios (illustrated in Fig. 3a), with sampling matching the empirical spatial and temporal distribution of our samples.

**Fig. 3.**
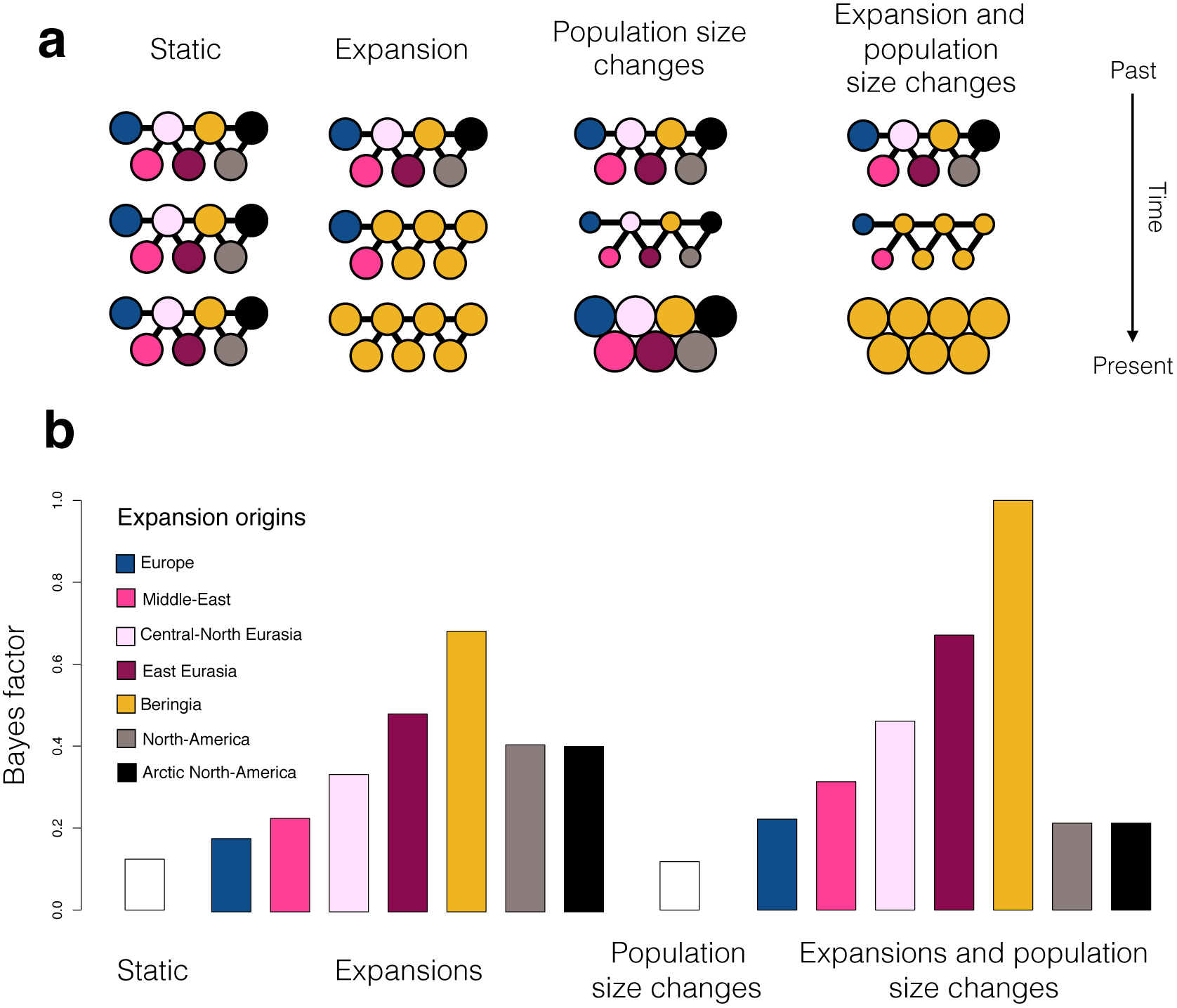
Spatially and temporally explicit analysis. (a) Illustration of the different scenarios, with circles representing one deme each for the seven different geographic regions (see panel b for colour legend and text for full description of the scenarios). Solid lines represent population connectivity. The *static* scenario (far left) shows stable populations through time. The *expansion* scenarios (middle left) shows how one deme (here yellow, corresponding to Beringia) expands and sequentially replaces the populations in all other demes (from top to bottom). The *population size change scenario* (middle right) illustrates how population size in the demes can change through time (large or small population size shown as large or small circles, respectively. We also show a combined scenario (far right) of both expansion and population size change. (b) Likelihood of each demographic scenario relative to the most likely scenario, shown as Bayes factors, estimated using Approximate Bayesian Computation analyses (see text for details). For expansion scenarios (including the combined expansion and population size changes), we colour code each bar according to the origin of the expansion (see colour legend).

The first scenario consisted of a constant population size and uniform movement between neighboring demes. This allowed us to test the null hypothesis that drift within a structured population alone can explain all the patterns observed in the mitochondrial tree. We then considered two additional demographic processes that could explain the observed patterns: 1) a temporal sequence of two population size changes that affected all demes simultaneously (thus allowing for a bottleneck); and 2) an expansion out of one of the demes, which had a continuous population through time and sequentially replaced the populations in the other demes (repeated for all seven possible expansion origins). We considered each demographic event in isolation as well as their combined effect (resulting in a total of 16 scenarios) and used Approximate Bayesian Computation (ABC) to calculate the likelihood of each scenario and estimate parameter values.

Both the null scenario and the scenario of only population size change in all demes were strongly rejected (Bayes Factor (BF) ≤ 0.1, Fig. 3b and Table S6), illustrating the power of combining a large dataset of ancient samples with statistical modeling. Scenarios that combined an expansion with a change in population size (bottleneck) were better supported than the corresponding scenarios (i.e. with the same origin) with constant population size (Fig. 3b). The best-supported scenario (BF 1, Fig. 4) was characterized by the combination of a rapid expansion of wolves out of the Beringian deme ~25,000 years ago (95% CI: 33,000-14,000 years ago) with a population bottleneck between 15,000 and 40,000 years ago, and limited gene flow between neighboring demes (see Table S7 and Fig. S15 for posterior distributions of all model parameters). We also found relatively strong support for a scenario that describes a wolf expansion out of the East Eurasian deme (BF 0.7) with nearly identical parameters to the best-supported scenario (Table S8 & Fig. S16). This can be explained by geographic proximity of East Eurasia to Beringia and the genetic similarity of wolves from these areas.

**Fig. 4.**
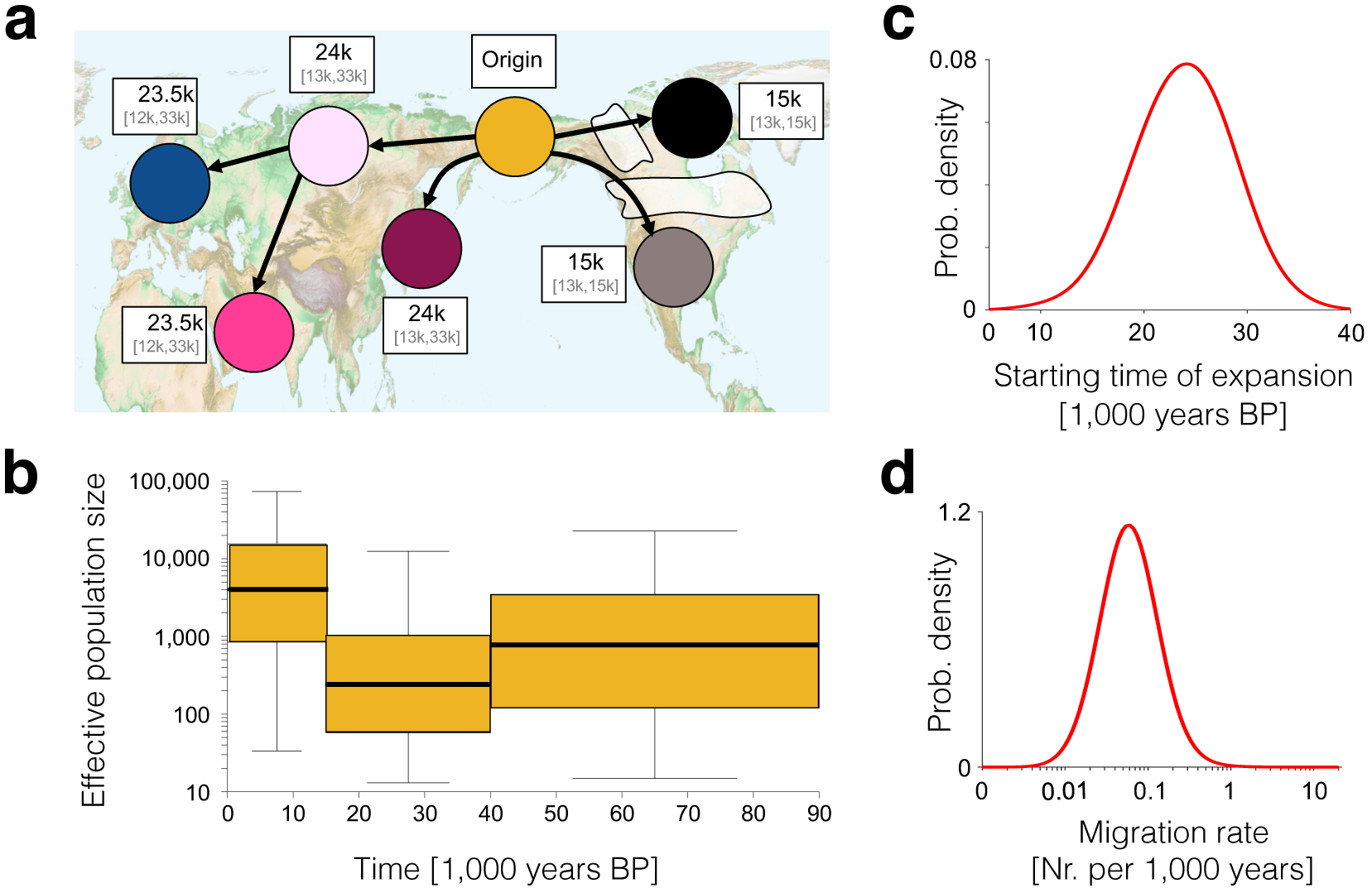
The inferred scenario of wolf demography from the Bayesian analysis using our spatially and temporally explicit model (see Fig. 3 and the main text). (a) Geographic representation of the expansion scenario (out of Beringia) with median and 95% CI for the date of the population replacement in each deme given in white boxes next to each deme. (b) Effective population size (thick line, boxes and whiskers show the median, interquartal range and 95% CI, respectively, for each time period). (c) Posterior distribution of migration rate and (d) starting time of expansion.

## Discussion

Recent whole-genome studies (3–5) found that modern grey wolves (*Canis lupus*) across Eurasia are descended from a single source population. The results of our analyses using both ancient and modern grey wolf samples (Fig. 1) within a spatially and temporally explicit modeling framework (Fig. 3), suggest that this process began ~25,000 (95% CI:13,000-33,000) years ago when a population of wolves from Beringia (or a Northeast Asian region in close geographic proximity) expanded outwards and replaced indigenous Pleistocene wolf populations across Eurasia (Fig. 4). The star-like topology of modern wolves observed in these whole genome studies is also consistent with our inferred scenario (Fig. 4) in which the wave of expansion is divided by geographic barriers, leading to divergence of subpopulations within the Northern Hemisphere due to subsequent limited gene flow.

In the Americas, the Beringian expansion was delayed due to the presence of ice sheets extending from Greenland to the northern Pacific Ocean (Fig. 4) (13). A recent study by (17) suggested that wolf populations that were extant south of these ice sheets were replaced by Eurasian wolves crossing the Beringian land bridge. Our analyses support the replacement of North American wolves (following the retreat of the ice sheets), and our more extensive ancient DNA sampling combined with a spatially explicit model has allowed us to narrow down the geographic origin of this expansion.

Were the wolves before and after this replacement ecologically equivalent? Analyses of wolf specimens have noted morphological differences between Late-Pleistocene and Holocene wolves: late Pleistocene specimens have been described as cranio-dentally more robust than the present-day grey wolves, as well as having specialized adaptations for carcass and bone processing (18–20) associated with megafaunal hunting and scavenging (21, 22). In contrast, the early Holocene archaeological record has only yielded a single sample with the Pleistocene wolf morphotype (in Alaska) (19), suggesting that this ecomorph had largely disappeared from the Northern Hemisphere by the Pleistocene-Holocene transition. This change in wolf morphology coincides with a shift in wolf isotope composition (23) and the disappearance of many megafaunal herbivores as well as other large predators, such as cave hyenas and cave lions, suggesting a possible change in the ecological niche of wolves.

It has been unclear whether the morphological change was the result of population replacement (genetic turnover), a plastic response to a dietary shift, or both. Our results suggest that the Pleistocene-Holocene transition was accompanied by a genetic turnover in most of the Northern Hemisphere wolf populations since most indigenous wolf populations experienced a large-scale replacement resulting in the loss of all native Pleistocene genetic lineages (Fig. 4). Similar population dynamics of discontinuity and replacement by conspecifics have been observed in several other large Pleistocene mammals in Europe including cave bears, woolly mammoths (24, 25), giant deer (24) and even humans (11, 26).

The geographic exception to this pattern of widespread replacement is Beringia, where we infer demographic continuity between late Pleistocene and Holocene wolf populations (Fig. 4). This finding is consistent with a recent study using the mtDNA control region by (27) that failed to reject continuity in this region, but at odds with a previous suggestion of genetic turnover in Beringia (19). This contradiction is likely the result of both the amount of data available and the analytical methodology: (19) used a short segment (427 bases long) of the mitochondrial control region and employed a descriptive phylogeographic approach, whereas our conclusions are based on an expanded dataset both in terms of sequence length, sample number, and geographic and temporal range (Fig. 1) and a formal hypothesis testing within a Bayesian framework (Figs. 3 and 4). As a consequence, the morphological and dietary shift observed in Beringian wolves between the late Pleistocene and Holocene (19) cannot be explained by a population turnover, but instead requires an alternative explanation such as adaptation or plastic responses to the substantial environmental and ecological changes that took place during this period. Indeed, grey wolves are a highly adaptable species. Studies of modern grey wolves have found that differences in habitat - specifically precipitation, temperature, vegetation, and prey specialization, can strongly affect their cranio-dental morphology (28–32).

The specific causal factors for the replacement of indigenous Eurasian wolves during the LGM by their Beringian conspecifics (and American wolves following the disappearance of the Cordilleran and Laurentide ice sheets) are beyond the scope of this study. However, one possible explanation may be related to the relatively stable climate of Beringia compared to the substantial climatic fluctuations that impacted the rest of Eurasia and Northern America during the late Pleistocene (33). These fluctuations have been associated with dramatic changes in food webs, leading to the loss of most of the large Pleistocene predators in the region (23, 34–36). In addition, the hunting of large Pleistocene predators by Upper Palaeolithic people (e.g. (37–39) may have also negatively impacted large carnivore populations (5). An interdisciplinary approach involving morphological, isotopic as well as genetic data is necessary to better understand the relationship between wolf population dynamics and dietary adaptations in the late Pleistocene and early Holocene period.

In summary, we have found that that, despite a continuous fossil record through the late Pleistocene, wolves experienced a complex demographic history involving population bottlenecks and replacements (Fig. 4). Our analysis suggests that long-range migration played an important role in the survival of wolves through the wave of megafaunal extinctions at the end of the last glaciation. These results will enable future studies to examine specific local climatic and ecological factors that enabled the Beringian wolf population to survive and expand across the Northern Hemisphere.

Lastly, the complex demographic history of Eurasian grey wolves reported here (Fig. 4) also has significant implications for identifying the geographic origin(s) of wolf domestication and the subsequent spread of dogs. For example, the limited understanding of the underlying wolf population structure may explain why previous studies have produced conflicting geographic and temporal scenarios. Numerous previous studies have focused on the patterns of genetic variation in modern domestic dogs, but have failed to consider potential genetic variation present in late Pleistocene wolf population, thereby implicitly assuming a homogenous wolf population source. As a result, both the domestication and the subsequent human-mediated movements of dogs were the only processes considered to have affected the observed genetic patterns in dog populations. However, both domestication from and admixture with a structured wolf population will have consequences for patterns of genetic variation within dogs.

In light of the complex demographic history of wolves (and the resulting population genetic structure) reconstructed by our analysis, several of the geographic patterns of haplotype distribution observed in previous studies, including differences in levels of diversity found within local dog populations (40), and the deep phylogenetic split between Eastern and Western Eurasian dogs (41), could have resulted from known admixture between domestic dogs and grey wolves (3, 5, 42, 43). Future analyses should therefore explicitly include the demographic history of wolves and demonstrate that the patterns of variation observed within dogs fall outside expectations that take admixture with geographically structured wolf populations into account.

## MATERIALS & METHODS

### Data preparation

We sequenced whole mitochondrial genomes of 40 ancient and 22 modern wolf samples. Sample information, including geographic locations, estimated ages and archaeological context information for the ancient samples, is provided in the Table S1 and Supplementary Information (SI) 1.2. Of the 40 ancient samples, 24 were directly radiocarbon dated for this study and calibrated using the IntCal13 calibration curve (see Table S1 for radiocarbon dates, calibrated age ranges and AMS laboratory reference numbers). DNA extraction, sequencing and quality filtering, and mapping protocols used are described in SI 2.

We included 16 previously published ancient mitochondrial wolf genomes (Table S1 and SI 2). In order to achieve a uniform dataset, we re-processed the raw reads from previously published samples using the same bioinformatics pipeline as for the newly generated sequences.

We subjected the aligned ancient sequences to strict quality criteria in terms of damage patterns and missing data (Figs. S3 – S5). First, we excluded all whole mitochondrial sequences that had more than 1/3 of the whole mitochondrial genome missing (excluding the mitochondrial control region – see below) at minimum three-fold coverage. Secondly, we excluded all ancient whole mitochondrial sequences that contained more than 0.1% of singletons showing signs of deamination damage typical for ancient DNA (C to T or A to G singletons). After quality filtering, we were left with 32 newly sequenced and 13 published ancient whole mitochondrial sequences (Table S1).

We also excluded sequences from archaeological specimens that postdate the end of Pleistocene and that have been identified as dogs (Table S1), since any significant population structure resulting from a lack of gene flow between dogs and wolves could violate the assumption of a single, randomly mating canid population. Some of the Pleistocene specimens used in the demographic analyses (TH5, TH12, TH14) have been argued to show features commonly found in modern dogs and have therefore been suggested to represent Paleolithic dogs (e.g. (22, 44–48). Here, we disregard such status calls because of the controversy that surrounds them ((49–52), and because early dogs would have been genetically similar to the local wolf populations form which they derived. This reasoning is supported by the close proximity of these samples to other wolf specimens confidently described as wolves in the phylogenetic tree (see Fig. S10).

Finally, we added 66 modern published wolf sequences from NCBI and two sequences from (3) (Table S1) resulting in a final dataset of 135 complete wolf mitochondrial genome sequences, of which 45 were ancient and 90 were modern. We used ClustalW alignment tool (version 2.1) (53) to generate a joint alignment of all genomes. In order to avoid the potentially confounding effect of recurrent mutations in the mitochondrial control region (54) in pairwise difference calculations, we removed this region from all subsequent analyses. This resulted in an alignment of sequences 15,466bp in length, of which 1301 sites (8.4%) were variable. The aligned dataset is located in Supplementary File S1.

### Phylogenetic analysis

We calculated the number of pairwise differences between all samples (Fig. S6) and generated a neighbor-joining tree based on pairwise differences (Fig. S7). This tree shows a clade consisting of samples exclusively from the Tibetan region and the Indian sub-continent that are deeply diverged from all ancient and other modern wolf samples (see also (55, 56)). A recent study of whole gnome data showed a complex history of South Eurasian wolves (5) that is beyond the scope of our study. While their neighbor-joining phylogeny grouped South Eurasian wolves with East and North East Asian wolves (Fig. 3 in Fan et al. (2016)), they cluster outside of all other grey wolves in a Principal Component Analysis (Fig. 4 in (5)), and also show a separate demographic history within a PSMC analysis (Fig. 5 in (5)). Because our study did not possess sufficient samples from the Himalayas and the Indian subcontinent to unravel their complex demography, we excluded samples from these regions and focused on the history of North Eurasian and North American wolves, for which we have good coverage through time and space.

We used PartitionFinder (57) and BEAST (v.1.8.0) (14) to build a tip calibrated wolf mitochondrial tree (with a strict global clock, see SI 3.2 for full details) from modern and directly dated ancient samples, and to estimate mutation rates for four different partitions of the wolf mitochondrial genome (see Tables S3 and S4 for results).

We used BEAST to molecularly date seven sequences from samples that were not directly radiocarbon dated (TH4, TH6, TH14, TU15) or that had been dated to a period beyond the limit of reliable radiocarbon dating (>48,000 years ago) (CGG12, CGG29, CGG32). We estimated the ages of the samples by performing a BEAST run where the mutation rate was fixed to the mean estimates from the previous BEAST analysis and all other parameter settings were set as described in the SI 3.2. We cross-validated this approach through a leave-one-out analysis where we sequentially removed a directly dated sample and estimated its date as described above. We find a close fit (R^2^=0.86) between radiocarbon and molecular dates (Fig. S9). We combined the seven undated samples with the 110 ancient and modern samples from the previous run and used a uniform prior ranging from 0 to 100,000 years to estimate the ages of the seven undated samples (see Table S5 for results).

Finally, in order to estimate the mitochondrial divergence time between the South Eurasian (Tibetan and Indian) and the rest of our wolf samples, we performed an additional BEAST run in which we included all modern and ancient grey wolves (*N* = 129) as well as five Tibetan and one Indian wolf, and used parameters identical to the ones described above. The age of the ancient samples was set as the mean of the calibrated radiocarbon date distribution (for radiocarbon dated samples) or as the mean of the age distribution from the BEAST analyses (for molecularly dated samples).

### Isolation by distance analysis

We performed isolation by distance (IBD) analyses to see the extent to which wolf mitochondrial genetic variation shows population structure. To this end, we regressed the pairwise geographic distances between 84 modern wolf samples (Table S1) against their pairwise genetic (mitochondrial) distances. The geographic distance between all sample pairs was calculated in kilometres as the great circle distance from geographic coordinates, using the Haversine Formula (58) to account for the curvature of the Earth as follows:

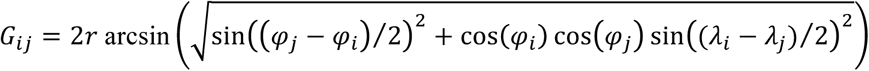

Where *G* is the distance in kilometers between individuals *i* and *j*; *ϕ_i_* and *ϕ_j_* are the latitude coordinates of individuals *i* and *j*, respectively; *λ_i_* and *λ_j_* are the longitude coordinates of individuals *i* and *j*, respectively; and r is the radius of the earth in kilometers. The pairwise genetic distances were calculated as the proportion of sites that differ between each pair of sequences (excluding the missing bases), using *dist.dna* function in the R package APE (59).

### Geographical deme definitions

We represented the wolf geographic range as seven demes, defined by major geographic barriers through time.

1. The *European* deme is bordered by open water from the North and the West (the Arctic and the Atlantic oceans, respectively); the Ural Mountains from the East; and the Mediterranean, the Black and the Caspian Sea and the Caucasus mountains from the South.
2. The *Middle-Eastern* deme consists of the Arabian Peninsula, Anatolia and Mesopotamia and is bordered by the Black Sea, the Caspian Sea and the Aral Sea in the North; the Indian Ocean in the South; the Tien Shen mountain range, the Tibetan Plateau and the Himalayas from the East; and the Mediterranean Sea in the West.
3. The *Central North Eurasian* deme consist of the Siberian Plateau and is bordered by the Arctic Ocean from the North; the Ural Mountains from the West; the Lena River and mountain ranges of North Eastern Siberia (Chersky and Verkhoyansk ranges) from the East; and the Tien Shen mountain range, the Tibetan Plateau and the Gobi Desert from South-East.
4. The *East Eurasian deme* is bordered by the Tien Shen mountain range, the Tibetan Plateau and Gobi desert from the West; the Pacific Ocean from the East; and the Lena river and the mountain ranges of North Eastern Siberia (Chersky and Verkhoyansk ranges) from the North.
5. The *Beringia* deme spans the Bering Strait, which was a land bridge during large parts of the Late Pleistocene and the Early Holocene. It is bordered to the West by the Lena River and mountain ranges of North Eastern Siberia (Chersky and Verkhoyansk ranges), and to the South and East by the extent of the Cordillerian and Laurentide ice sheets during the Last Glacial Maximum.
6. The *Arctic North America* deme consists of an area of the North American continent east of the Rocky Mountains and west of Greenland, that was covered by ice during the last Glaciation and is at present known as the Canadian Arctic Archipelago.
7. The *North America* deme consists of an area in the Northern American sub-continent that was south of the Cordillerian and Laurentide ice sheets during the last glaciation (13).

### Demographic scenarios

We tested a total of 16 demographic scenario combinations, from four different kinds of demographic scenarios (illustrated in Fig. 3a in the main text):

1) Static model (the null hypothesis) – neighboring demes exchange migrants, no demographic changes.
2) Bottleneck scenarios – demes exchange migrants as in the static model but populations have different size in different time periods. We consider three time periods: 0-15k years ago, 15k-40k years ago, and >40k years ago.
3) Expansion scenarios - demes exchange migrants like in the static model but a single deme experiences an expansion starting between 5k and 40k years ago (at a minimum rate of 1,000 years per deme, so the whole world could be colonized within 3,000 years or faster).
4) Combinations of scenarios 2 & 3.

### Population genetic coalescent framework

We implemented coalescent population genetic models for the different demographic scenarios to sample gene genealogies.

In the static scenario, we simulated local coalescent processes (60) within each deme (scaled to rate 1/*K* per pair of lineages, where *K* is the mean time to most recent common ancestor in a deme and is thus proportional to the effective population size). In addition, we moved lineages between demes according to a Poisson process with rate *m* per lineage. To match the geographic and temporal distribution of the data, we represented each sample with a lineage from the corresponding deme and date.

The bottleneck scenario was implemented as the static one but with piecewise constant values for *K* as a function of time. We considered three time periods, each with its own value of *K* (*K*_1_, *K*_2_ and *K*_3_), motivated by the archaeological and genetic evidence of wolf population changes described in the main text. The first time period was from present to early Holocene, 0-15k years ago. The second time period extended from early Holocene to late Pleistocene and covered the last glacial maximum, 15-40k years ago. Finally, the third time period covered the late Pleistocene and beyond, i.e. 40k years ago and older.

The population expansion scenarios were based on the static model but with an added population expansion model with founder effects and replacement of local populations (we refer to populations not yet replaced by the expansion as “indigenous”). Starting at time *T*, the population expanded from the initial deme and replaced its neighboring populations. After the start of the expansion, the expansion proceeded in fixed steps of *ΔT* (in time). At each step, colonized populations replaced neighboring indigenous populations (if an indigenous deme bordered to more than one colonized deme, these demes contributed equally to the colonization of the indigenous deme). In the coalescent framework (that simulates gene genealogies backwards in time) the colonization events corresponds to forced migrations from the indigenous deme to the source deme. If there were more than one source deme, the source of each lineage was chosen randomly with equal probability. Finally, founder effects during the colonization of an indigenous deme were implemented as a local, instantaneous population bottleneck in the deme (after the expansion), with a severity scaled to give a fixed probability *x* of a coalescent event for each pair of lineages in the deme during the bottleneck (61). (*x*=1 correspond to a complete loss of genetic diversity in the bottleneck, and *x*=0 corresponds to no reduction in genetic diversity.)

Finally, the combined scenario of population expansion and bottlenecks was implemented by making the population size parameter *K* in the population expansion model time dependent as in the population bottleneck model.

### Approximate Bayesian Computation analysis

We used Approximate Bayesian Computation (ABC) analysis (62) with ABCtoolbox (63) to formally test the fit of our different demographic models. This approach allows formal hypothesis testing using likelihood ratios in the cases where the demographic scenarios are too complex for a direct calculation of the likelihoods given the models. We used the most likely tree from BEAST (see SI 3.2 for details) as data, and simulated trees using the coalescent simulations described above.

To match the assumption of random mixing within each deme in the population genetic model, we removed closely related sequences if they came from the same geographic location and time period, by randomly retaining one of the closely related sequences to be included in the analysis (Table S1, column “Samples_used_in_Simulation_Analysis”).

To robustly measure differences between simulated and observed trees we use the matrix of time to most recent common ancestor (TMRCA) for all pairs of samples. This matrix also captures other allele frequency based quantities frequently used as summary statistics with ABC, such as F_ST_, as they can be calculated from the components of this matrix.

In principle the full matrix could be used, but in practice it is necessary to use a small number of summary statistics for ABC to work properly (63). To this end, we grouped our seven demes into four super demes (Fig. S14), based on geographic proximity and genetic similarity in the dataset, and used mean TMRCA within each super deme and mean TMRCAs between all super demes as summary statistics in the ABC analysis. The four super demes are 1) Europe; 2) Middle East; 3) North East Eurasia, Beringia and East Eurasia combined; and 4) Artic and Continental North America combined. This resulted in 10 summery statistics in total.

An initial round of fitting the model showed that all scenarios underestimate the within-super deme TMRCA for the Middle East, while the rest of the summary statistics were well captured by the best fitting demographic scenarios. This could be explained by a scenario where the Middle East was less affected by the reduction in population size during the last glacial maximum. However, we currently lack sufficient number of samples from this area to explicitly test a more complex scenario such as this hypothesis. To avoid outliers biasing the likelihood calculations in ABC (63) we removed this summary statistic, resulting in nine summary statistics in total.

For each of the 16 scenarios we performed 1 billion simulations with randomly chosen parameter combinations, chosen from the following parameter intervals for the different scenarios:

- The static scenario: *m* in [0.001,20] and *K* in [0.01,100].
- The bottleneck scenarios: *m* in [0.001,20] and *K*_1_, *K*_2_, *K*_3_ in [0.01,100].
- The expansion scenarios: *m* in [0.001,20], *K* in [0.01,100], *x* in [0,1], *T* in [5,40] and *ΔT* in [0.001,1]. For expansion out of the North American scenario and the expansion out of the Arctic North American scenario, the glaciation and during the LGM in North American and sea level rise during the de-glaciation mean that T must be in the range [9,16]
- The combined bottleneck and expansion scenarios: *m* in [0.001,20], *K*_1_,*K*_2_,*K*_3_ in [0.01,100], *x* in [0,1], *T* in [5,40] and *ΔT* in [0.001,1].

The parameter *m* is measured in units of 1/1,000 years, and *T*, *ΔT*, *K*, *K_1_*, *K_2_* and *K*_3_ are measured in units of 1,000 years. The parameters *x, T* and *ΔT* were sampled according to a uniform distribution over the interval, while all other parameters were sampled from a uniform distribution of their log-transformed values. To identify good parameter combinations for ABC, we first calculated the Euclidian square distances between predicted and observed statistics and restricted analysis to parameter combinations within the lowest tenth distance percentile. We then ran the ABCtoolbox (63) on the accepted parameter combinations to estimate posterior distributions of the model parameters, and to calculate the likelihood of each scenario as described in the ABCtoolbox manual.

See Table S6 for ABC likelihoods and Bayes factors for all demographic scenarios tested.

See Tables S7 and S8 for posterior probability estimates and Figs. S.15 and S16 for posterior density distributions for estimated parameters (*ΔT*, *T*, log_10_ *K*_1_, log_10_ *K*_2_, log_10_ *K*_3_, log_10_ *m*, *x*) in the two most likely models (An expansion out of Beringia with a population size change and an expansion out of East Eurasia with a population size change).

### Data availability

New sequences are available to download from GenBank database (accession numbers XX-XX). The raw sequence reads are available from ENA database (accession numbers XX-XX). The scripts used in the analyses are available up on request from L.L. and A.E.

## Acknowledgements

The authors are grateful to Daniel Klingberg Johansson & Kristian Murphy Gregersen from the Natural History Museum of Denmark; Gabriella Hürlimann from the Zurich Zoo; Jane Hopper from the Howlett’s & the Port Lympne Wild Animal Parks; Cyrintha Barwise-Joubert & Paul Vercammen from the Breeding Centre for Endangered Arabian Wildlife; Link Olson from the University of Alaska Museum of the North; Joseph Cook & Mariel Campbell from the Museum of Southwestern Biology; Lindsey Carmichael & David Coltman from the University of Alberta; North American Fur Auctions; Department of Environment Nunavut and Environment and Natural Resources Northwest Territories for DNA samples from the modern wolves.

The authors are also grateful to the staff at the Danish National High-Throughput Sequencing Centre for technical assistance in the data generation; the Qimmeq project, funded by The Velux Foundations and Aage og Johanne Louis-Hansens Fond, for providing financial support for sequencing ancient Siberian wolf samples; the Rock Foundation (New York, USA) for supporting radiocarbon dating of ancient samples from the Yana site;to Stephan Nylinder from the Swedish Museum of Natural History for advice on phylogenetic analyses and Terry Brown from the University of Manchester for comments on this manuscript. L.L., K.D. & G.L. were supported by Natural Environment Research Council, UK (grant numbers NE/K005243/1, NE/K003259/1); LL. was also supported by the European Research Council grant (339941-ADAPT); A.M. & A.E. were supported by the European Research Council Consolidator grant (grant number 647787-LocalAdaptation); L.F. & G.L. were supported by the European Research Council grant (ERC-2013-StG 337574-UNDEAD); T.G was supported by European Research Council Consolidator grant (681396-Extinction Genomics) & Lundbeck Foundation grant (R52-5062); O.T. was supported by the National Science Center, Poland (2015/19/P/NZ7/03971) with funding from EU’s Horizon 2020 program under the Marie Skłodowska-Curie grant agreement (665778) and Synthesys Project (BETAF 3062); V.P., E.P. & P.N. were supported by the Russian Science Foundation grant (N16-18-10265 RNF); A.P. was supported by the Max Planck Society; M.L-G. was supported by Czech Science Foundation grant (GAČR15-06446S).

## Author contributions

L.L., O.T., M.T.P.G., J.K., G.L., A.E. and A.M. designed the research; O.T., M-H.S.S., V.J.S., K.E.W., M.S.V., I.K.C.L., N.W. and G.S. performed ancient DNA laboratory work with input from J.K., M.T.P.G., H.S., K-H.H., R.S.M. and K-H.H.; M-H.S.S. performed modern DNA laboratory work with input from M.T.P.G; O.T., J.A.S.C. and L.L. performed bioinformatic analyses; L.L., A.E. and A.M. designed the population genetic analyses; L.L. Performed phylogenetic analyses; A.E. implemented the spatial analyses framework; L.L and A.E. performed spatial analyses; M.G., J.B., V.V.P., E.Y.P., P.A.N., S.E.F., J.E-L., A.W.K., B.G., H.N., H-P.U. and M.L-G. provided samples; V.V.P., M.G., M. L-G., H.B., H.N., A.W.K., E.Y.P. and P.A.N. provided context for archaeological samples; A.P., M.G., H.B. and K.D. Helped setting the results of genetic analyses into an archaeological context; A.M., M.T.P.G., A.J.H., G.L., J.K., E.W. and K.D. secured funding for the project; L.L., O.T. and A.E. wrote the initial draft of the manuscript with input from A.M.; L.L., O.T. and A.E wrote the manuscript and the supplementary information with input from A.P., M.G., H.B., M-H.S.S., M.T.P.G., K.E.W., A.M., G.L and K.D.; V.J.S., L.F., A.W.K., K-H.H., A.J.H., R.S.M., H.S., G.S., V.V.P., E.Y.P., P.A.N. and J.E-L. provided comments to the manuscript and/or to the supplementary information.

## References

1. Puzachenko AY, Markova AK (2016) Diversity dynamics of large- and medium-sized mammals in the Late Pleistocene and the Holocene on the East European Plain: Systems approach. Quat Int. doi:10.1016/j.quaint.2015.07.031.

2. Sotnikova M, Rook L (2010) Dispersal of the Canini (Mammalia, Canidae: Caninae) across Eurasia during the Late Miocene to Early Pleistocene. Quat Int 212(2):86–97.

3. Freedman AH, et al. (2014) Genome Sequencing Highlights the Dynamic Early History of Dogs. PLOS Genet 10(1):e1004016.

4. Skoglund P, Ersmark E, Palkopoulou E, Dalén L (2015) Ancient Wolf Genome Reveals an Early Divergence of Domestic Dog Ancestors and Admixture into High-Latitude Breeds. Curr Biol 25(11):1515–1519.

5. Fan Z, et al. (2016) Worldwide patterns of genomic variation and admixture in gray wolves. Genome Res 26(2):163–173.

6. Larson G, et al. (2012) Rethinking dog domestication by integrating genetics, archeology, and biogeography. Proc Natl Acad Sci 109(23):8878–8883.

7. Groucutt HS, et al. (2015) Rethinking the dispersal of Homo sapiens out of Africa. Evol Anthropol Issues News Rev 24(4):149–164.

8. Rambaut A (2000) Estimating the rate of molecular evolution: incorporating non-contemporaneous sequences into maximum likelihood phylogenies. Bioinformatics 16(4):395–399.

9. Drummond AJ, Nicholls GK, Rodrigo AG, Solomon W (2002) Estimating Mutation Parameters, Population History and Genealogy Simultaneously From Temporally Spaced Sequence Data. Genetics 161(3):1307–1320.

10. Rieux A, et al. (2014) Improved calibration of the human mitochondrial clock using ancient genomes. Mol Biol Evol:msu222.

11. Posth C, et al. (2016) Pleistocene Mitochondrial Genomes Suggest a Single Major Dispersal of Non-Africans and a Late Glacial Population Turnover in Europe. Curr Biol 26(6):827–833.

12. Warmuth V, et al. (2012) Reconstructing the origin and spread of horse domestication in the Eurasian steppe. Proc Natl Acad Sci 109(21):8202–8206.

13. Raghavan M, et al. (2015) Genomic evidence for the Pleistocene and recent population history of Native Americans. Science:aab3884.

14. Drummond AJ, Suchard MA, Xie D, Rambaut A (2012) Bayesian Phylogenetics with BEAUti and the BEAST 1.7. Mol Biol Evol 29(8):1969–1973.

15. Thalmann O, et al. (2013) Complete Mitochondrial Genomes of Ancient Canids Suggest a European Origin of Domestic Dogs. Science 342(6160):871–874.

16. Nielsen R, Beaumont MA (2009) Statistical inferences in phylogeography. Mol Ecol 18(6):1034–1047.

17. Koblmüller S, et al. (2016) Whole mitochondrial genomes illuminate ancient intercontinental dispersals of grey wolves (Canis lupus). J Biogeogr 43(9):1728–1738.

18. Kuzmina I., Sablin MV (1993) Pozdnepleistotsenovyi pesets verhnei Desny. Materiali po mezozoickoi i kainozoickoi istorii nazemnykh pozvonochnykh (Trudy 17 Zoologicheskogo Instituta RAN 249), pp 93–104.

19. Leonard JA, et al. (2007) Megafaunal Extinctions and the Disappearance of a Specialized Wolf Ecomorph. Curr Biol 17(13):1146–1150.

20. Baryshnikov GF, Mol D, Tikhonov AN (2009) Finding of the Late Pleistocene carnivores in Taimyr Peninsula (Russia, Siberia) with paleoecological context. Russ J Theriol 8(2):107–113.

21. Fox-Dobbs K, Leonard JA, Koch PL (2008) Pleistocene megafauna from eastern Beringia: Paleoecological and paleoenvironmental interpretations of stable carbon and nitrogen isotope and radiocarbon records. Palaeogeogr Palaeoclimatol Palaeoecol 261(1):30–46.

22. Germonpré M, et al. (2017) Palaeolithic and prehistoric dogs and Pleistocene wolves from Yakutia: Identification of isolated skulls. J Archaeol Sci 78:1–19.

23. Bocherens H (2015) Isotopic tracking of large carnivore palaeoecology in the mammoth steppe. Quat Sci Rev 117:42–71.

24. Stuart AJ, Kosintsev PA, Higham TFG, Lister AM (2004) Pleistocene to Holocene extinction dynamics in giant deer and woolly mammoth. Nature 431(7009):684–689.

25. Palkopoulou E, et al. (2013) Holarctic genetic structure and range dynamics in the woolly mammoth. Proc R Soc B 280(1770):20131910.

26. Fu Q, et al. (2016) The genetic history of Ice Age Europe. Nature 534(7606):200–205.

27. Ersmark E, et al. (2016) From the Past to the Present: Wolf Phylogeography and Demographic History Based on the Mitochondrial Control Region. Front Ecol Evol 4. doi:10.3389/fevo.2016.00134.

28. Geffen E, Anderson MJ, Wayne RK (2004) Climate and habitat barriers to dispersal in the highly mobile grey wolf. Mol Ecol 13(8):2481–2490.

29. Pilot M, et al. (2006) Ecological factors influence population genetic structure of European grey wolves. Mol Ecol 15(14):4533–4553.

30. O’Keefe FR, Meachen J, Fet EV, Brannick A (2013) Ecological determinants of clinal morphological variation in the cranium of the North American gray wolf. J Mammal 94(6):1223–1236.

31. Flower LOH, Schreve DC (2014) An investigation of palaeodietary variability in European Pleistocene canids. Quat Sci Rev 96(Supplement C):188–203.

32. Leonard JA (2015) Ecology drives evolution in grey wolves. Evol Ecol Res 16(6):461–473.

33. Clark PU, et al. (2012) Global climate evolution during the last deglaciation. Proc Natl Acad Sci 109(19):E1134–E1142.

34. Lister AM, Stuart AJ (2008) The impact of climate change on large mammal distribution and extinction: Evidence from the last glacial/interglacial transition. Comptes Rendus Geosci 340(9–10):615–620.

35. Hofreiter M, Stewart J (2009) Ecological Change, Range Fluctuations and Population Dynamics during the Pleistocene. Curr Biol 19(14):R584–R594.

36. Lorenzen ED, et al. (2011) Species-specific responses of Late Quaternary megafauna to climate and humans. Nature 479(7373):359–364.

37. Münzel SC, Conard NJ (2004) Change and continuity in subsistence during the Middle and Upper Palaeolithic in the Ach Valley of Swabia (south-west Germany). Int J Osteoarchaeol 14(3–4):225–243.

38. Germonpré M, Hämäläinen R (2007) Fossil Bear Bones in the Belgian Upper Paleolithic: The Possibility of a Proto Bear-Ceremonialism. Arct Anthropol 44(2):1–30.

39. Cueto M, Camarós E, Castaños P, Ontañón R, Arias P (2016) Under the Skin of a Lion: Unique Evidence of Upper Paleolithic Exploitation and Use of Cave Lion (Panthera spelaea) from the Lower Gallery of La Garma (Spain). PLOS ONE 11(10):e0163591.

40. Wang G-D, et al. (2016) Out of southern East Asia: the natural history of domestic dogs across the world. Cell Res 26(1):21–33.

41. Frantz LAF, et al. (2016) Genomic and archaeological evidence suggest a dual origin of domestic dogs. Science 352(6290):1228.

42. Verardi A, Lucchini V, Randi E (2006) Detecting introgressive hybridization between free-ranging domestic dogs and wild wolves (Canis lupus) by admixture linkage disequilibrium analysis. Mol Ecol 15(10):2845–2855.

43. Godinho R, et al. (2011) Genetic evidence for multiple events of hybridization between wolves and domestic dogs in the Iberian Peninsula. Mol Ecol 20(24):5154–5166.

44. Sablin M, Khlopachev G (2002) The Earliest Ice Age Dogs: Evidence from Eliseevichi 1. Curr Anthropol 43(5):795–799.

45. Germonpré M, et al. (2009) Fossil dogs and wolves from Palaeolithic sites in Belgium, the Ukraine and Russia: osteometry, ancient DNA and stable isotopes. J Archaeol Sci 36(2):473–490.

46. Druzhkova AS, et al. (2013) Ancient DNA Analysis Affirms the Canid from Altai as a Primitive Dog. PLOS ONE 8(3):e57754.

47. Germonpré M, Lázničková-Galetová M, Sablin MV (2012) Palaeolithic dog skulls at the Gravettian Předmostí site, the Czech Republic. J Archaeol Sci 39(1):184–202.

48. Germonpré M, Lázničková-Galetová M, Losey RJ, Räikkönen J, Sablin MV (2015) Large canids at the Gravettian Předmostí site, the Czech Republic: The mandible. Quat Int 359–360:261–279.

49. Crockford SJ, Kuzmin YV (2012) Comments on Germonpré et al., Journal of Archaeological Science 36, 2009 “Fossil dogs and wolves from Palaeolithic sites in Belgium, the Ukraine and Russia: osteometry, ancient DNA and stable isotopes”, and Germonpré, Lázkičková-Galetová, and Sablin, Journal of Archaeological Science 39, 2012 “Palaeolithic dog skulls at the Gravettian Předmostí site, the Czech Republic.” J Archaeol Sci 39(8):2797–2801.

50. Morey DF (2014) In search of Paleolithic dogs: a quest with mixed results. J Archaeol Sci 52:300–307.

51. Drake AG, Coquerelle M, Colombeau G (2015) 3D morphometric analysis of fossil canid skulls contradicts the suggested domestication of dogs during the late Paleolithic. Sci Rep 5:8299.

52. Perri A (2016) A wolf in dog’s clothing: Initial dog domestication and Pleistocene wolf variation. J Archaeol Sci 68:1–4.

53. Larkin MA, et al. (2007) Clustal W and Clustal X version 2.0. Bioinformatics 23(21):2947–2948.

54. Excoffier L, Yang Z (1999) Substitution rate variation among sites in mitochondrial hypervariable region I of humans and chimpanzees. Mol Biol Evol 16(10):1357–1368.

55. Sharma DK, Maldonado JE, Jhala YV, Fleischer RC (2004) Ancient wolf lineages in India. Proc R Soc Lond B Biol Sci 271(Suppl 3):S1–S4.

56. Aggarwal RK, Kivisild T, Ramadevi J, Singh L (2007) Mitochondrial DNA coding region sequences support the phylogenetic distinction of two Indian wolf species. J Zool Syst Evol Res 45(2):163–172.

57. Lanfear R, Calcott B, Ho SYW, Guindon S (2012) PartitionFinder: Combined Selection of Partitioning Schemes and Substitution Models for Phylogenetic Analyses. Mol Biol Evol 29(6):1695–1701.

58. Sinnott R (1984) Virtues of the Haversine. Sky Telesc 68(2):159.

59. Paradis E, Claude J, Strimmer K (2004) APE: Analyses of Phylogenetics and Evolution in R language. Bioinformatics 20(2):289–290.

60. Kingman JFC (1982) The coalescent. Stoch Process Their Appl 13(3):235–248.

61. Eriksson A, Mehlig B (2004) Gene-history correlation and population structure. Phys Biol 1(4):220.

62. Beaumont MA, Zhang W, Balding DJ (2002) Approximate Bayesian Computation in Population Genetics. Genetics 162(4):2025–2035.

63. Wegmann D, Leuenberger C, Neuenschwander S, Excoffier L (2010) ABCtoolbox: a versatile toolkit for approximate Bayesian computations. BMC Bioinformatics 11(1):116.

